# Efficacy and safety of a novel vaginal medical device in recurrent bacterial vaginosis: an international multicentre clinical trial

**DOI:** 10.1101/674705

**Authors:** Filippo Murina, Ciprian Crişan, Marius Biriş, Daniela Sîrbu, Dionisio Franco Barattini, Luca Ivan Ardolino, Elena Casolati

## Abstract

Several risk factors have been identified but the etiology and pathogenesis of Bacterial vaginosis (BV) are still not completely understood, and the recurrence rate of BV remains high despite adequate chemotherapy treatment.

The primary objective of the study was to assess the effectiveness of a new vaginal medical device, which contains polycarbophil, 0.04 % lauryl glucoside, and glycerides (Polybactum^®^ – Effik Italia), in reducing BV recurrence rate.

This was a multicenter, open label, not comparative study performed in Italy and Romania. Female subjects over 18-years-old affected by recurrent BV were included. The latest episode was diagnosed by Amsel criteria 6-9 days before the start of the study and treated with vaginal metronidazole (gel 0.75% mg for 5 days or ovules 500 mg for 7 days). The recurrence was defined by at least 2 episodes in the previous 12 months. Polybactum^®^ vaginal ovules, day 1-4-7, were started within the 12th and the 24th hr after the end of metronidazole therapy and repeated monthly for 3 cycles.

The first 41 patients enrolled were evaluated for an interim analysis 6 months after the study started; 2 patients interrupted the trial, leaving 39 evaluable subjects. The recurrence rate was significantly reduced compared to previous published data (10.26% vs 40% p<0.001). In 35 patients without recurrence, the assessment of Lactobacillus vaginal flora performed by phase contrast microscopy evidenced a significant improvement form baseline (p=0.022) The Investigator global assessment of tolerability was excellent in 38 out of 39 cases.

**IMPORTANCE:** Bacterial vaginosis (BV) is the most common vaginal disorder in women of childbearing age. In BV, Lactobacillus species, which are predominant in a healthy vaginal flora, are replaced by anaerobes, mainly Gardnerella vaginalis. BV is responsible for more than 60% of vulvovaginal infections and has been linked to serious, potentially life-threatening conditions, including: pelvic inflammatory disease, postoperative infections, acquisition and transmission of the human immunodeficiency virus, preterm birth, and several adverse pregnancy outcomes. Our research showed that 3 monthly cycles of Polybactum^®^ ovules administered after one course of metronidazole vaginal therapy can reduce the rate of Bacterial vaginosis recurrence and improve the vaginal milieu, favouring the growth of vaginal lactobacillus species. Taken together our results confirm that Polibactum^®^ is a safe and effective treatment to reduce BV recurrence rate after a first line therapy with metronidazole.

## INTRODUCTION

Bacterial vaginosis (BV) is the most common vaginal disorder in women of childbearing age (1). In BV, Lactobacillus species, which are predominant in a healthy vaginal flora, are replaced by anaerobes, mainly *Gardnerella vaginalis*, but also *Atopobium vaginae*, *Mobiluncus mulieris*, *Prevotella bivia*, and *Fusobacterium nucleatum*.

BV is responsible for more than 60% of vulvovaginal infections and has been linked to serious, potentially life-threatening conditions, including: pelvic inflammatory disease, postoperative infections, acquisition and transmission of the human immunodeficiency virus, preterm birth, and several adverse pregnancy outcomes (2, 3, 4). Hydrogen peroxide and lactic acid–producing lactobacilli generally dominate the healthy vaginal epithelium, acting as a protective surfactant layer. This leads to an acidic pH that inhibits the adhesion and the growth of other bacteria, including opportunistic pathogens on the vaginal epithelium (5). During the development of BV, the normal vaginal microbiota composition changes, characterized by a decrease in the number of such lactobacilli species and an increase in the number of several pathogenic bacteria, mainly anaerobes (6). Although at least 50% of the infected women have no symptoms, the most common complaints are of abnormal vaginal discharge and fishy odor. Usually, the vulvovaginal examination normal, apart from a discharge described as watery and gray. A correct diagnosis relies on finding three out of four Amsel criteria (abnormal gray discharge, high vaginal pH, positive amine test, and greater than 20% clue cells on saline microscopy), with a sensitivity of 92%(7). Metronidazole (oral or topical), tinidazole (oral), and clindamycin (oral or topical) are all recommended as initial treatments (8). The etiology and pathogenesis of this disease are not completely understood, because of the various BV-determining risk factors, and the treatment is not always effective, resulting in high recurrence rates. After treatment, up to 58% of women might have a single recurrence or more within 12 months. Limited data is available regarding optimal management strategies for women with persistent or recurrent BV (8).

Recurrence are relates to the inability to offer a long-term defensive barrier, thus facilitating relapses. Furthermore, the resistance of pathogens to multiple drugs is a (relatively new) health problem that needs alternative treatments to be developed.

Recent studies have found that 90% of women with and 10% without BV have a complex polymicrobial biofilm, which can be demonstrated by electron microscopy of vaginal biopsies. On women with this disease, the biofilm consists primarily of *Gardnerella vaginalis*, sometimes including *Atopobium vaginae* (9). With standard antibiotic regimens, the bacterial load may decrease, but the biofilm may not be eliminated, thus setting the stage for recurrence after treatment. Slime produced by bacteria is a mechanism of forming biofilm that shields and protects microbes against the effects of antibiotics.

The objective of this study was to assess the effectiveness and safety of a new vaginal product that contains polycarbophil, 0.04 % lauryl glucoside, and glycerides (Polybactum^®^ – Effik Italia) in reducing the rate of BV recurrence.

The rationale relates to the specific bacteriostatic action of Polybactum^®^, which inhibits bacterial growth, and its mucoadhesive property impairing the formation of biofilm produced by *Gardnerella vaginalis* and other bacteria. Furthermore, the product ensures an acidifying effect on the vaginal pH which favors the growth of lactobacillus microbiota and at the same time maintains a hostile environment for the recolonization of the vagina by the polymicrobial flora involved in BV.

## MATERIALS AND METHODS

### Trial design

This was a multicenter, open label, not comparative study performed in Italy and Romania. The study protocol was identified with the acronym POLARIS (Polybactum^®^ to assess Recurrent Bacterial Vaginosis) and was approved by the National Agency for Medicines and Medical Devices in Romania (Agentia Nationala a Medicamentului si a Dispozitivelor Medicale), notified to the Italian Ministry of Health and approved by the local Independent Ethics Committees pertaining to the investigational sites. The study was registered in clinicaltrials.gov as NCT02863536 (with attached full protocol).

During the trial, the protocol was amended to allow an interim analysis of 6 months after the start of patient enrolment.

### Participants

Female subjects over 18-years-old affected by recurrent BV were included. They were diagnosed based on the Amsel criteria (10) in the 6-9 days before baseline and treated with metronidazole vaginal formulations (gel for 5 days or ovules for 7 days). The recurrence was defined by at least 2 episodes of (BV) in the last 12 months, including the BV episode treated before baseline. Additional inclusion criteria: informed consent form (ICF) signed before starting the trial and the status of non-lactating women or lactating, but not amenorrhoeic women.

The exclusion criteria: pregnancy; candidiasis or mixed vaginitis; HIV or another immunodeficiency; known allergy to metronidazole or to Polybactum^®^ ingredients; prostitution; ongoing menstruation or pre-menopause/menopause; patients concomitantly included in different interventional clinical trials; unwillingness to provide the informed consent to the trial; time between the last day of last menses and baseline visit >16 days or ≤5 days (to avoid bias in case of a menstrual bleeding occurring during the first cycle with the tested medical device and the consequent need to interrupt its administration); participation in another clinical trial during the last month.

The sites were: Vittore Buzzi Hospital (Milan, coordinator site), AIED Center in Rome (Italy) and 3 private clinics specialized in gynaecology located in Timisoara (Romania). In each center, a lead Investigator was appointed to be responsible for the identification, recruitment, data collection, and completion of Case Report Forms, (CRFs) for patient adherence to protocol and for the sample collections to be transported to the local laboratory. The Investigators were specialists in gynaecology and trained in Good Clinical Practice (GCP). During the study, all the patients presented spontaneously at the Investigator’s visit. No advertising was used to increase the enrolment rate.

### Interventions

Prior to any study procedure, each patient was informed about the nature and purpose of the trial, the benefits and the risks, and was asked to sign an ICF. The tested medical device administration (Polybactum^®^ vaginal ovule) started within the 12th and the 24th hr after the end of metronidazole vaginal treatment (5 g of 0.75% gel once daily for 5 days or 500 mg ovules once daily for 7 days) and continued for 3 cycles of treatment (minimum 72 and maximum 84 days); the duration of each cycle was one week, with the tested medical device administered as follows: one ovule inserted in the vagina on day 1, one ovule on day 4 and the last ovule on day 7. On the baseline visit, each patient received a total number of 9 ovules for the whole study duration (3 ovules for 3 cycles). The Investigator advised the patient to lay in a supine position for a couple minutes after the ovule had been inserted.

The baseline visit and the first cycle of Polybactum^®^ fell within the 6th and the 16th day after the menstrual bleeding and after the end of metronidazole treatment. In the second and the third cycle, Polybactum^®^ was administered immediately after the end of the previous menstrual bleeding.

In any case, the Investigator could always decide to stop administering the medical device for safety purposes or to prescribe other therapies if considered necessary for the patient’s health.

The Sponsor, Effik Italia, supplied the investigational product. Patients were reminded to return all investigational product packages to the Investigator. At the end of the study, the Contract Research Organization (CRO) personnel involved in the trial performed the accountability of the tested medical device.

During the trial, there were disallowed: vaginal tampons; use of an etonogestrel/ethinyl estradiol vaginal ring (Nuvaring^®^) or an intrauterine device; oral or vaginal antibiotic therapy or other vaginal therapies (like douching, spermicide); oral or vaginal probiotics (e.g. vaginal lactobacilli); other products or medication to treat BV.

### Primary and secondary outcomes

The primary outcome was the recurrence of BV identified by Amsel criteria (10), determined at baseline and at final visit; a positive diagnosis of BV required meeting three of the following four criteria:

- vaginal pH greater than pH 4.5;
- proportion of clue cells ≥20% of total epithelial cells in the vaginal fluid;
- presence of white and thin vaginal discharge;
- fishy smell at whiff test.

BV was excluded if only two or less criteria were found at baseline. On the other hand, the patient was evaluated as a treatment failure when at least two criteria were met at the final visit. The primary outcome was measured at baseline and the final visit for all patients.

The secondary outcomes were the following: vaginal Lactobacillus microbiota assessed by an optical microscopy at baseline and at the final visit to evaluate the vaginal microflora return rate to normality (11) after Polybactum^®^ treatment; signs and symptoms of BV (vaginal discharge, burning, erythema, dyspareunia). Vaginal discharge over the last 24 hrs was evaluated using this analogic 3-point scale: 0= not present or physiological in quantity, colour, and type; 1 = mild abnormal (abnormal quantity with normal colour and type); 2 = abnormal quantity, colour and type.

Burning intensity over the last 24 hrs was evaluated using an analogic 5-point scale: 0 = not present; 1 = mild; 2 = moderate; 3 = severe; 4 = unbearable. Grade of erythema was evaluated using the analogic 5-point scale: 0 = no symptoms; 1 = slight; 2 = moderate; 3 = marked; 4 = very marked. Dyspareunia was evaluated by a dichotomic scale: 0 = absent; 1 = present. All secondary outcomes were measured at baseline and the final visit considering value change from baseline. Secondary outcome also included the patient assessing the efficacy at the last visit using the 4-point scale: 1 = very good improvement; 2 = good improvement; 3 = moderate improvement and 4 = negligible improvement. Safety was evaluated collecting and analysing the adverse events during the study period and by a global assessment of safety performed by any Investigator using an analogic 4-point scale: 1 = excellent, 2 = good, 3 = fair and 4 = poor.

The study schedule included a visit at baseline (day 0), and a final visit (day 72 to day 84). In addition, the Investigators planned to have 3 phone contacts with the patient: the first at day 28±1 after the last day of last menses, and the following two after 28±1 day apart. On any phone contact the Investigator checked if the patient had any of the BV symptoms and, in this case, performed an unscheduled visit to verify the BV recurrence based on Amsel criteria.

### Sample size determination and statistical methods

According to published data, the mean recurrence rate of BV after a first episode is from 30 to 50% within 3 months after a first efficacious medical therapy (12); in the present study patients were not allowed oral or vaginal antibiotic therapy after metronidazole treatment for the 3 month study duration; therefore it should be realistic to have a 40% as mean recurrence rates in this study as well. In addition, in the recently collected data on recurrence rates post-treatment with Polybactum^®^ (Effik Italia SpA, unpublished data), the correlation between paired observation was 2% and after applying continuity correction, the study would require a sample size of 44 pairs to achieve a power of 80% and a one-sided significance of 5% for detecting a difference of 0.25 between marginal proportions. Considering the drop-out rate, 55 were enrolled (one group chi square test than a proportion equals user specified value non-inferiority). The level of significance of <0.5 was considered statistically significant at 95% confidence interval. If a subject was missing information for one or more variables, the missing data were not replaced. If a subject were involved in violation of inclusion/exclusion criteria, the respective data were excluded from the analysis. Safety was evaluated in terms of adverse event findings. All subjects receiving at least one dose of treatment were included in the safety analysis. Analysis comparing the values between visits was done using t-test or chi squared test for quantitative variables, McNemar test for binary variables and symmetry test for qualitative variables. Kaplan Meier curves were used to analyse the time-to-event regarding BV recurrences. Statistical analysis was performed using SAS 9.2 (SAS Institute Inc., Cary, NC, USA).

### Guidelines and legislation

The trial was performed in accordance with UNI EN ISO 14155:2012, the ethical principles of the current version of the Declaration of Helsinki (64th WMA General Assembly, Fortaleza, Brazil, October 2013) and the Directive 91/507/EEC, Guidelines for GCP. The privacy of patient data was protected following the current policy in Italy and in Romania.

## RESULTS

The data presented are an interim analysis performed 6 months after the first patient enrolled. Participants were recruited in 3 Romanian sites from September 8th, 2016 until December 13th, 2016. As shown in the flow diagram in Figure 1, out of 41 enrolled patients, 2 were excluded from PP analysis because they interrupted the trial for personal reasons not related to safety and 39 subjects were considered evaluable. One patient of these 39 did not complete the study period for intercurrent recurrence.

**FIG 1:**
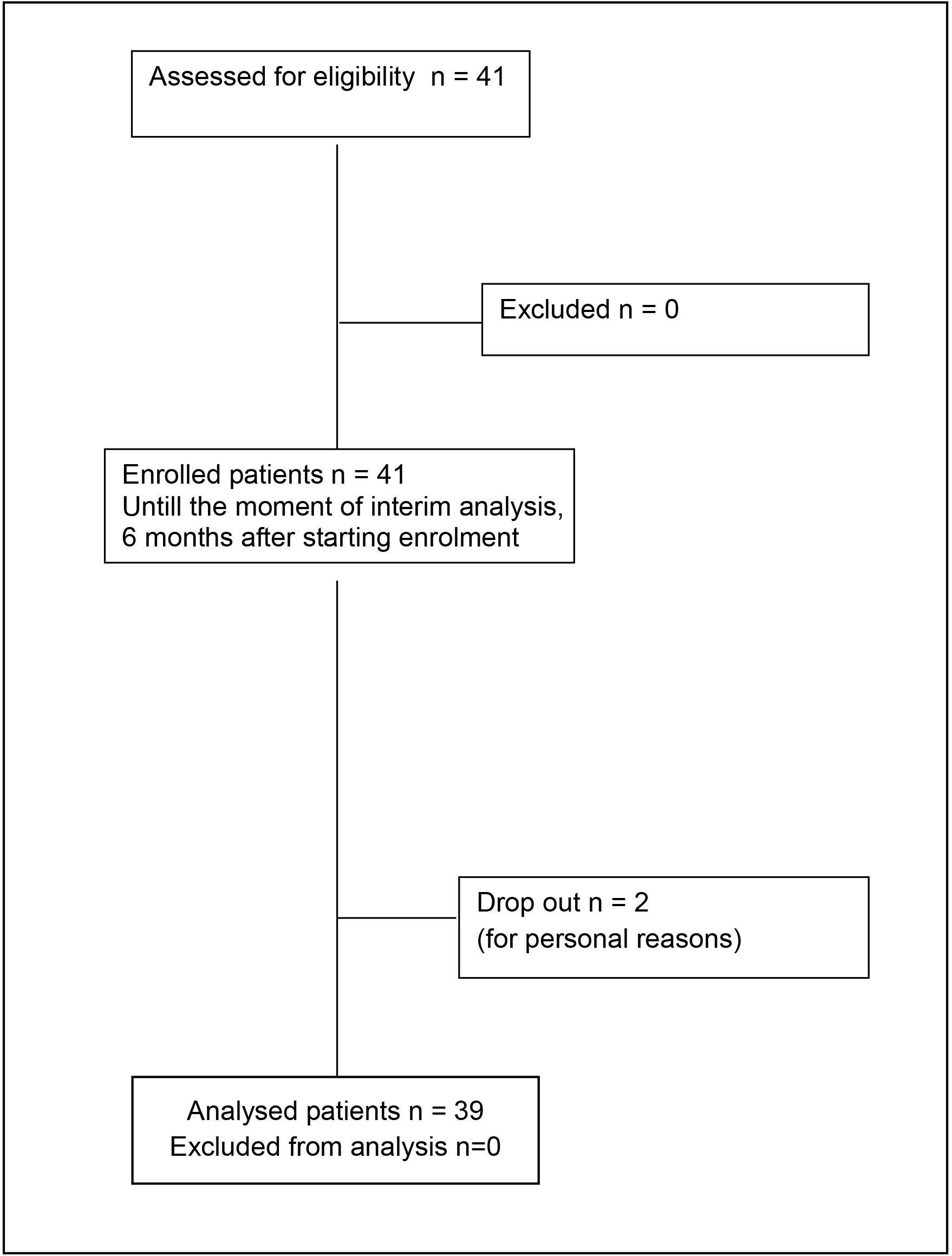
Flow diagram of the study

Age group distribution is summarized in Table 1. Treatment compliance rate (the total number of administered ovules from the total number of given ovules) was 97.01% for all the patients enrolled, and 100% considering only the patients who completed the trial. No relevant findings were evidenced in the medical history and during the physical examination. All the patients completed an antibiotic treatment with metronidazole vaginal formulation (ovules for 7 days) and no concomitant medication was recorded during the screening evaluation.

**TABLE 1.**
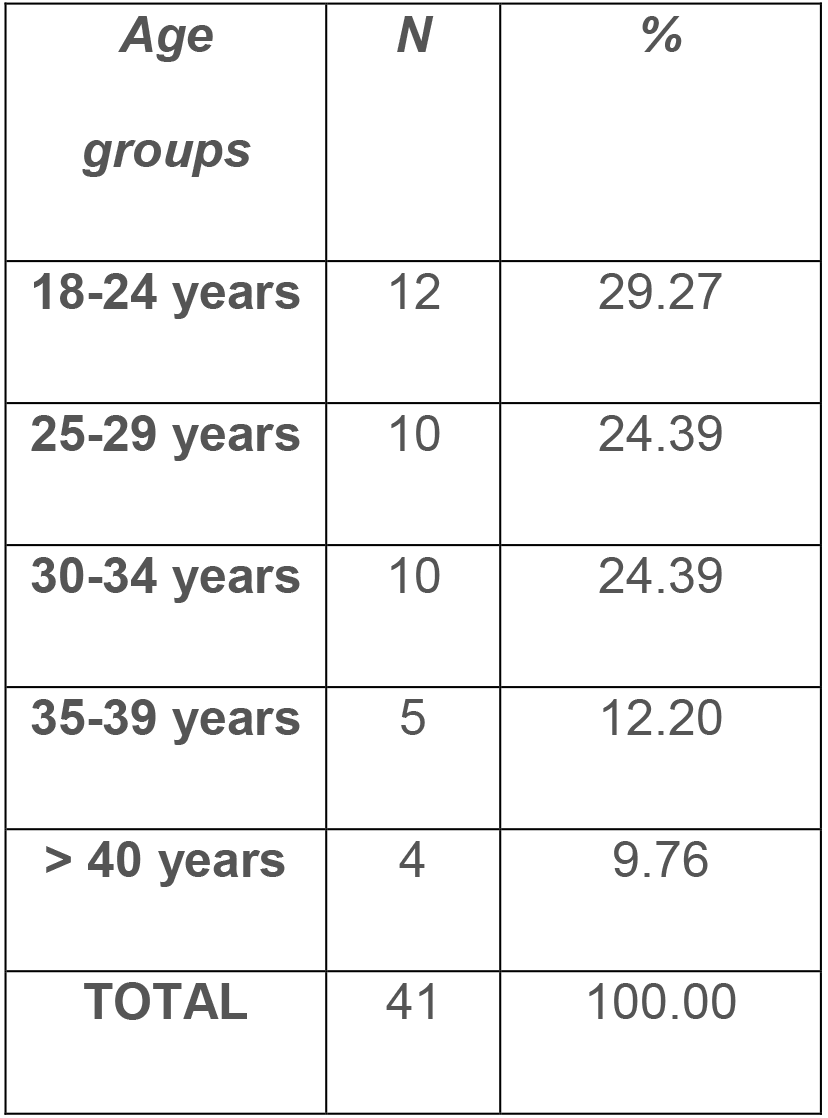
Subjects group distribution by age

| ***Age***<br>***groups*** | ***N*** | ***%*** |
| --- | --- | --- |
| **18-24 years** | 12 | 29.27 |
| **25-29 years** | 10 | 24.39 |
| **30-34 years** | 10 | 24.39 |
| **35-39 years** | 5 | 12.20 |
| **> 40 years** | 4 | 9.76 |
| **TOTAL** | 41 | 100.00 |

One of the inclusion criteria stated that only subjects with two or more episodes of BV in the last 12 months should enter the clinical trial. As observed in Table 2, 26 subjects (63.41%) had 2 episodes of BV in the last 12 months, with 12 subjects (29.27%) having 3 episodes of BV, and 3 subjects (7.32%) with 4 episodes of BV.

**TABLE 2.** Number of patients with multiple BV episodes in the last 12 months by trial site

|  | **Site 4** | **Site 6** | **Site 7** | **TOTAL**<br>**(%)** |
| --- | --- | --- | --- | --- |
| **2 episodes** | 0 | 25 | 1 | 26<br>(63.41) |
| **3 episodes** | 5 | 6 | 1 | 12<br>(29.27) |
| **4 episodes** | 1 | 2 | 0 | 3<br>(7.32) |
| **TOTAL** | 6 | 33 | 2 | 41<br>(100.00) |

In the analysis of the primary objective of the study, at the final visit (day 72 to day 84 from baseline), 4 recurrences were identified. Therefore, the recurrence rate of BV after 3 months of treatment with Polybactum^®^ was 10.26% (4 cases in 39). A statistically significant difference (p < 0.001) was evidenced comparing this value with recurrence rate data (40%) published in the medical literature (12). To compare the actual recurrences belonging to our data with those from medical literature, we carried out a time-to-event analysis, also called Kaplan-Meier survival curves. Data are censored (the event did not occur during clinical trial period or subject’s drop-out) or uncensored (the event occurred during the clinical trial). As observed from Figure 2, after 1 month of treatment, the probability of not having a BV recurrence is 97.6%; the same probability, at the final visit (after 3 months of treatment with Polybactum^®^) is 89.7%. Therefore, we can conclude that the true recurrence rate of BV after 3 months of treatment with Polybactum^®^ is significantly smaller than the 40% reported by medical literature.

**FIG 2:**
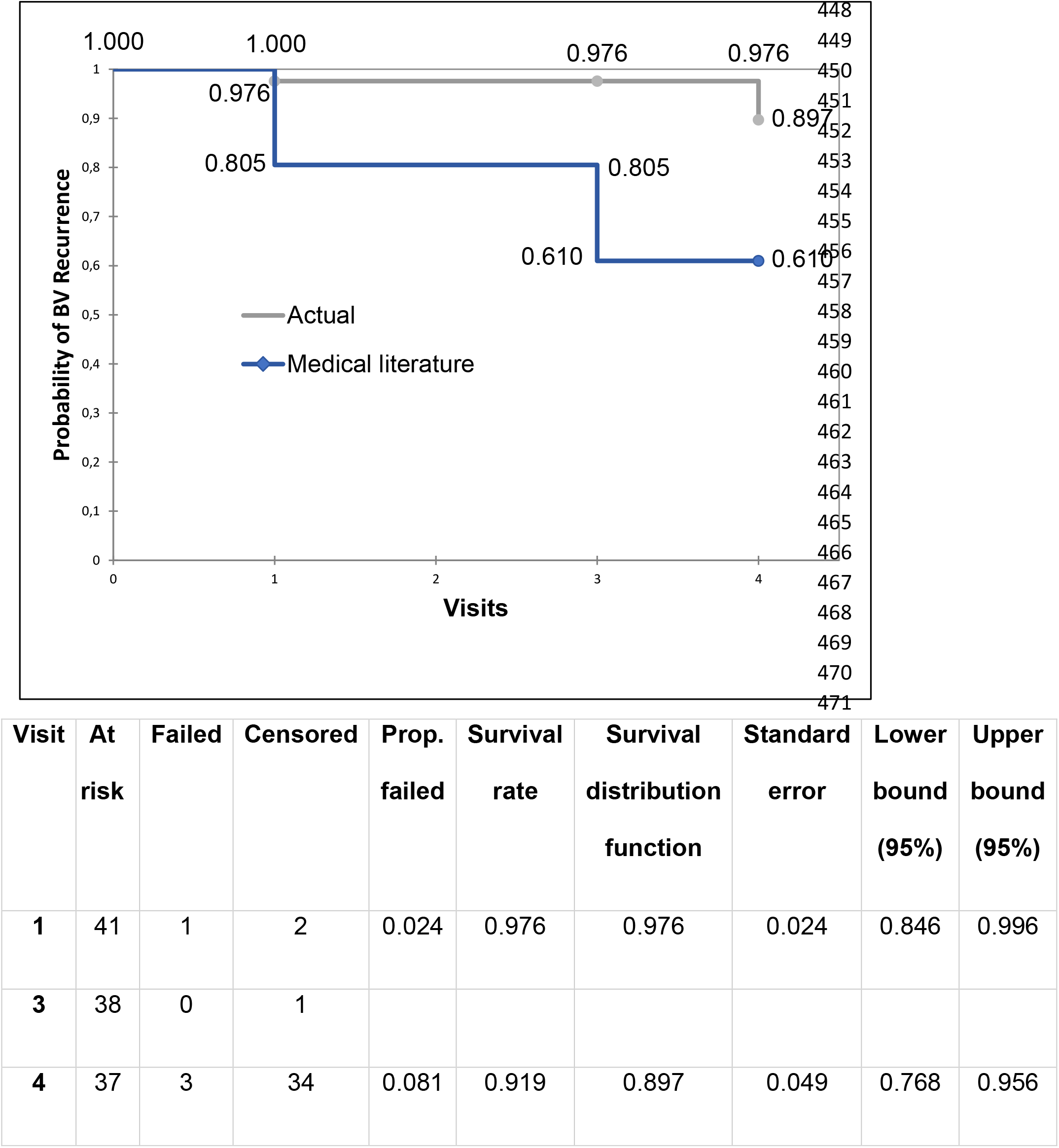
Kaplan-Meier survival curve for Bacterial Vaginosis recurrence event

The evaluation of Vaginal Lactobacillus microbiota by microscopy using phase contrast of vaginal secretions was measured from lowest to highest concentration by the following 5-point scale: absent, 1+, 2+, 3+, 4+ (13) at the baseline visit, and at the final visit (after 3 months of Polybactum^®^ treatment). For 26 out of 39 patients (66.67%) the treatment had a beneficial or neutral effect on the Lactobacillus concentration levels and only in 13 patients (33.33%) the concentration levels worsened (Figure 3). Chi-squared test evidenced that the differences between these values were at the limit of statistical significance (p-value = 0.054). However, analysing the 35 patients without recurrences only, a statistically significant difference (Maxwell-Stuart asymptotic homogeneity test, p-value = 0.022) was evidenced in Lactobacillus concentration levels between baseline and final visit. (Figure 4) Throughout the study period, symptoms associated with BV (vaginal discharge, burning, erythema and dyspareunia) were monitored and recorded by the Principal Investigators in the CRFs and by subjects themselves in the Patient’s Diary. The baseline and final data are reported in Table 3. It could be emphasized that the patients with abnormal symptoms at final visit are the four patients with recurrences and 1 additional patient with isolated mild vaginal discharge. These extremely positive results were confirmed by the global assessments of efficacy performed at the end of the study by patients. In fact, it was rated as very good per patient’s assessments for 87.18% of them, good for 5.13%, moderate for 2.56%, and negligible for 5.13% of the patients.

**FIG 3:**
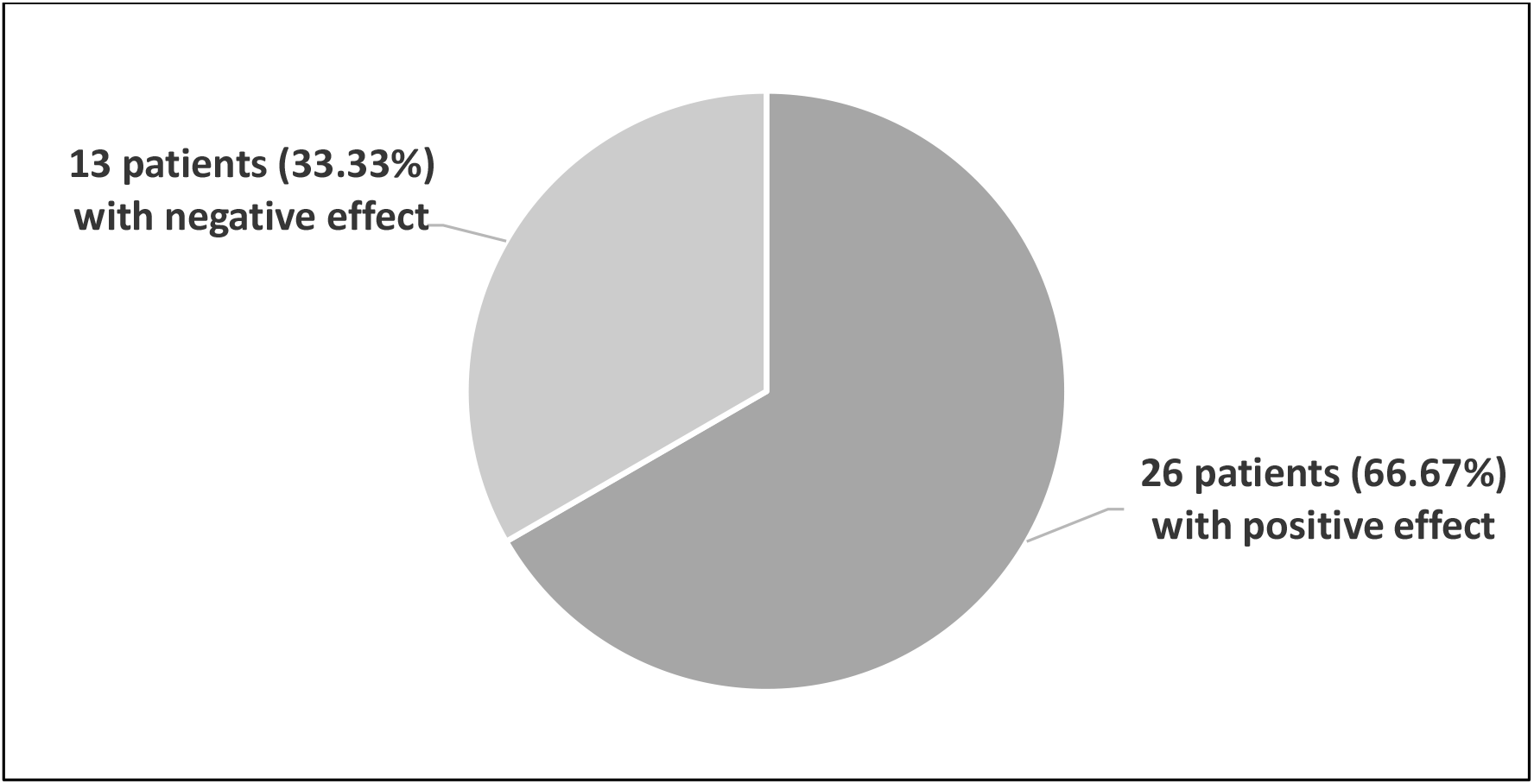
The effect on Lactobacillus concentration level change between baseline and final visit

**FIG 4:**
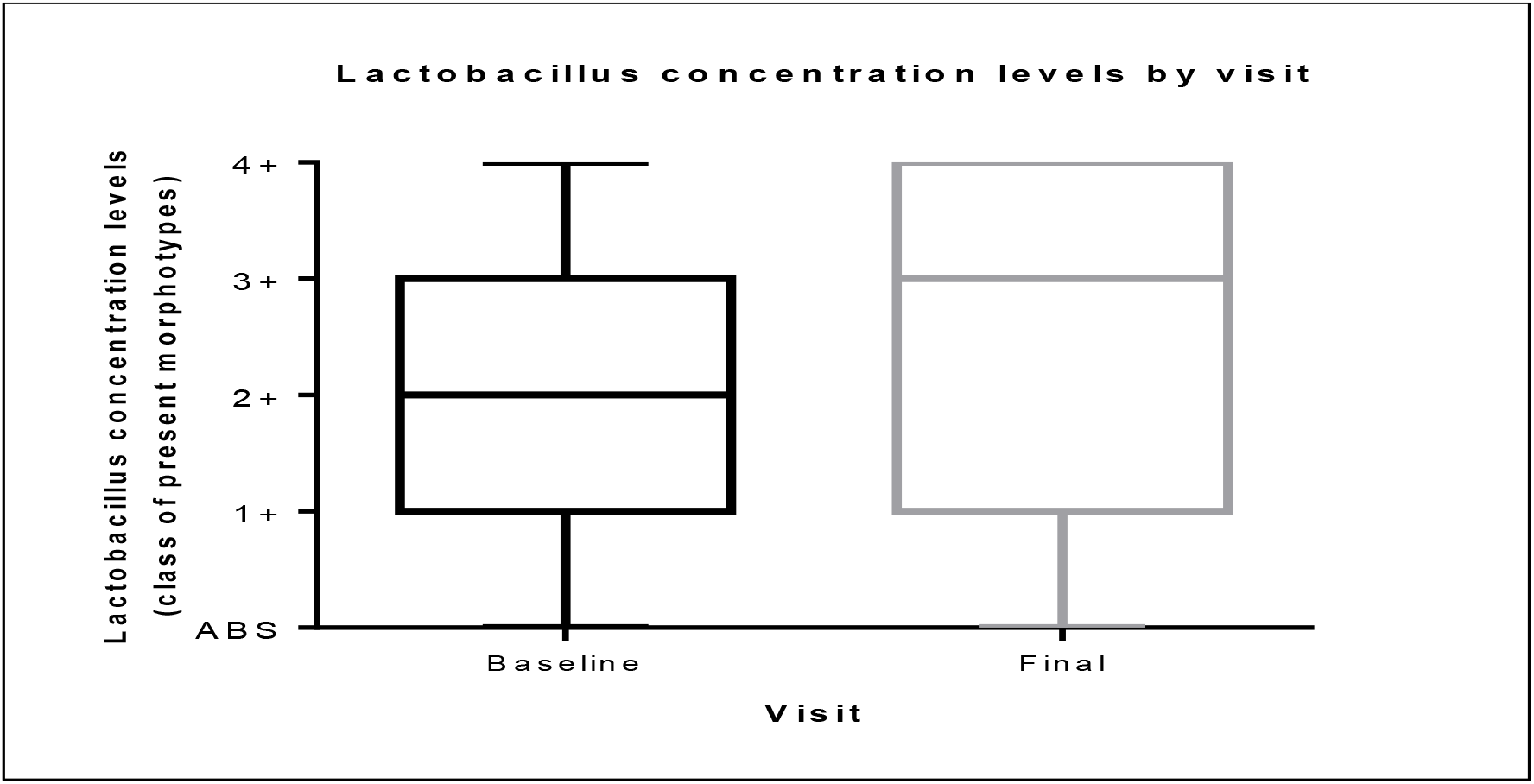
Lactobacillus concentration values between baseline and final visit on the 35 patients without recurrences (Maxwell-Stuart asymptotic marginal homogeneity test, p-value = 0.022)

**TABLE 3.** Number of symptoms and severity at different visits/phone contacts

|  | ***Vaginal***<br>***Discharge*** | ***Burning*** | ***Erythema*** | ***Dyspareunia*** |
| --- | --- | --- | --- | --- |
| ***Baseline Visit*** |  |  | 2 Slight |  |
| ***1<sup>st</sup> Phone***<br>***Contact***<br>***(1 Month)*** | 4 Mild<br>1 Abnormal | 1 Moderate<br>1 Unbearable | 1 Moderate<br>1 Marked<br>1 Very Marked | 3 Present |
| ***2<sup>nd</sup> Phone***<br>***Contact***<br>***(2 Months)*** | 1 Mild |  |  | 1 Present |
| ***3<sup>rd</sup> Phone***<br>***Contact***<br>***(3 Months)*** | 6 Mild | 4 Moderate | 4 Moderate | 4 Present |
| ***Final Visit***<br>***(3-5 days after***<br>***last Phone***<br>***Contact)*** | 4 Mild<br>1 Abnormal | 3 Moderate<br>1 Severe | 1 Slight<br>2 Moderate<br>1 Very Marked | 4 Present |
| ***Number of***<br>***Symptoms per***<br>***Visits*** | 17 | 10 | 13 | 12 |

Vaginal discharge: 0= not present; 1 =mild abnormal; 2= abnormal.
Burning: 0 = not present; 1= mild; 2 = moderate; 3 = severe; 4 = unbearable.
Erythema: 0 = no symptoms; 1 = slight; 2 = moderate; 3 = marked; 4= very marked.
Dyspareunia: 0= absent; 1 = present.

During the study, two adverse events have been reported. The cases (mild local itching and mild viral respiratory infection) were evaluated as not related to the tested medical device, recovered after a few days and in both cases the Investigator ruled out the suspicion of BV Recurrence. After the symptoms, the patient who experienced itching dropped out from the study for personal reasons, having completed only 1 cycle of treatment.

The Investigator’s global assessment of tolerability was excellent (38 out of 39 cases) for 97.43% and good for 2.57% (1 out of 39) of the patients.

## DISCUSSION

According to literature data, given the high prevalence of BV, there is an urgent need to develop products that effectively treat the condition and prevent its recurrence. In our study, the recurrence rate of BV after 3 months of treatment with Polybactum^®^ was 10.26%, significantly lower than the 40% presented in the medical literature. It is consented that BV involves the presence of a dense, structured and polymicrobial biofilm, primarily constituted by *Gardnerella vaginalis* clusters, strongly adhered to the vaginal epithelium.

Since the bacteria within biofilms are not effectively eliminated by the immune system or fully destroyed by antibiotics, biofilm-related infections tend to persist and thus, unsurprisingly, BV tends to have a high rate of relapse and recurrence (5).

Although the biofilm was shown to contain high concentrations of a variety of bacterial groups, *Gardnerella vaginalis* was the predominant constituent and it is now accepted that biofilms in BV are strongly associated with *Gardnerella vaginalis* (14). It was shown that *Gardnerella vaginalis* was able to adhere to and displace precoated protective lactobacilli from the vaginal epithelial cells, while other BV-associated anaerobes, such as *Atopobium vaginae* and *Prevotella*, were less virulent (15). Consequently, the current paradigm is that the establishment of a *Gardnerella vaginalis* biofilm is a required event for initiation and progression of BV (16).

The new product tested in our study contains polycarbophil, 0.04% lauryl glucoside, glycerides; all constituents that have important effects on the main mechanism involved in the pathogenesis of BV. In fact, lauryl glucoside has a specific bacteriostatic action which inhibits *Gardnerella vaginalis* growth. It was proven that *Gardnerella vaginalis* growth is reduced for 48 hrs by the contact with Polybactum^®^ (already at 24 hrs) (17). The growth of inhibitory activity of the new vaginal product was also demonstrated against *Streptococcus agalactiae* and *Neisseria gonorrhoeae*. Polybactum^®^ main components are polycarbophil (a safe film forming agent well known for its lack of toxicity) and lauryl glucoside, a non-ionic surfactant reinforcing the film-forming effect by reducing surface tension. The combination of these factors gives the product the unique characteristics of a mucoadhesive property impairing the formation of the biofilm produced by *Gardnerella vaginalis*. Finally, an acidifying effect was observed on the vaginal pH which favors the growth of lactobacillus microbiota maintaining a hostile environment for the polymicrobial flora involved in BV to recolonize the vagina. Therefore, vaginal biofilms play a key role not only in BV pathogenesis, but also in its treatment failure and recurrence. Most proposed non-antibiotic therapies for BV are vaginal or oral probiotics, with the aim to restore the normal vaginal microbiota. Several trials evaluated the use of combined metronidazole and/or clindamycin therapy and vaginal probiotics to prevent BV recurrence, but large, randomized, placebo-controlled trials with standardized outcomes are needed to confirm the efficacy of this therapeutic approach for BV (18).

Our study focused on demonstrating that the use of Polybactum^®^ in the treatment of BV not only reduces the rate of relapses, but it also improves the microbiological parameters. In fact, for 26 out of 35 patients without recurrences (74.28%), the clinical trial had a beneficial or neutral effect on the Lactobacillus concentration levels. From a clinical point of view, since metronidazole (the most commonly prescribed antibiotic to treat BV) causes only transient suppression of the *Gardnerella vaginalis* populations in the vagina (19), the combined therapy with this antibiotic plus Polybactum^®^ can successfully reduce the anaerobic bacterial pathogens responsible for BV, delaying or preventing the relapse of vaginal infection, thus preventing the administration of antibiotics in repetitive courses.

Our study has certain limitations, such as the lack of a placebo treatment arm and only a 3-month follow-up after the end of metronidazole treatment. This last problem will be corrected by the amendment already submitted to the EC to allow a follow up period of 12 months after the end of the antibiotic therapy.

In any case, this study strengthens the evidence supporting the use of specific new vaginal products with well demonstrated activity associated with the creation and maintenance of a vaginal biofilm that hinders the persistence of an infection caused by BV.

## ACKNOWLEDGEMENTS

Effik Italia (http://www.effikitalia.it/), the study Sponsor, offered a grant support. Effik Italia had no role in the study design, data collection and interpretation, or on the decision to submit the work for publication.

Warm thanks to Ramona Petrita and Andreea-Denisa Toma for support in data management and to Marius Ardelean for statistical analysis.

FM, CC, MB, and DS declare no conflict of interest. DFB is employed at Opera CRO, the Contract Research Organization that managed the study, EC is a medical consultant for Effik Italia, LIA is employed at Effik Italia.

## Authors’ contribution

FM originally conceived the project; FM and DFB draw up the study design and protocol. The final manuscript was written and approved by all authors.

